# Improved MALDI-TOF MS based antimicrobial resistance prediction through hierarchical stratification

**DOI:** 10.1101/2022.04.13.488198

**Authors:** Caroline Weis, Bastian Rieck, Sebastian Balzer, Aline Cuénod, Adrian Egli, Karsten Borgwardt

**Affiliations:** Machine Learning and Computational Biology Lab, D-BSSE, ETH Zurich, Switzerland; SIB Swiss Institute of Bioinformatics, Switzerland; Applied Microbiology Research, Department of Biomedicine, University of Basel, Basel, Switzerland; Division of Clinical Bacteriology and Mycology, University Hospital Basel, Basel, Switzerland

## Abstract

**Motivation:** Predicting antimicrobial resistance using MALDI-TOF mass spectrometry based machine learning is a fast-growing field of research. Recent advances in machine learning methods specifically designed for MALDI-TOF mass spectra have outperformed established classification approaches. However, classification performance was observed to have a large standard deviation between different train–test splits. We hypothesise that this variance is caused by the underlying phylogenetic structure between microbial samples, which is implicitly reflected in their MALDI-TOF MS profiles, but not taken into account during the training of a model.

**Results:** In this paper, we propose to infer this structure from the dataset—using agglomerative hierarchical clustering—and consider it during the dataset splitting between train and test. We show that incorporating such phylogenetic structure into the antimicrobial resistance prediction scenario leads to an improved classification performance. Average precision was increased from 42.3 to 47.1 for ciprofloxacin resistance prediction in *Escherichia coli* and from 44.6 to 50.8 for amoxicillin-clavulanic acid resistance prediction in *Staphylococcus aureus* using a Gaussian process classifier with a MALDI-TOF MS specific kernel. We envision that these results will support the quick and reliable identification of antimicrobial resistances, thus increasing patient well-being and reducing healthcare costs.

**Availability:** All data is available for download and code available as an easy-to-use Python package under https://github.com/BorgwardtLab/maldi_PIKE at branch `maldi_stratification`.

**Supplementary information:** Supplementary information at the end of document.

## 1 Introduction

Matrix-assisted laser desorption ionisation time-of-flight (MALDI-TOF) mass spectrometry (MS) is a fast and high-throughput method for the characterisation of bacteria due to its ability to analyse complex peptide mixtures. During the last few years, it has become a widely-used method for microbial species identification [De Bruyne et al., 2011]. The measurements of MALDI-TOF MS are represented in a set of mass spectra, known to be highly specific for different microbes. Coupled with antimicrobial resistance profiles, i.e. *a priori* knowledge about whether a given microbial is susceptible to a certain antimicrobial drug, MALDI-TOF MS was shown to yield accurate and confident predictions. Specifically, a large-scale study of MALDI-TOF based machine learning demonstrated accurate predictions and the potential for clinical utility [Weis et al., 2022] and a novel kernel method based on diffusion processes was recently shown to outperform existing methods [Weis et al., 2020a]. The improved performance was marred by high standard deviations—different train–test splits exhibited highly-varying predictive performance values. We conjecture that these differences are caused by the underlying phylogenetic structure of samples; the genetic information to infer the phylogeny is commonly unavailable in datasets. The train–test splits used to optimise parameters of machine learning algorithms should ideally follow the structure of the complete dataset, which is the best approximation we have of the *true* structure within the data. While the experimental setup of Weis et al. [2020a] ensures that the train–test split is stratified in terms of resistance information, it does *not* account for phylogenetic structure which could be distributed differently in the train and test datasets. We hypothesise that MALDI-TOF mass spectra contain this information in an *implicit* fashion, making a more refined description necessary for stratification. Although the ground truth phylogenetic tree cannot be reproduced, as the dataset consists of only a few thousand samples per species and only MALDI-TOF MS data, i.e. no additional genetic data, is available, we assume that phylogenetic structure can be partially inferred using similarity-based clustering. Ideally, each cluster should represent a different phylogenetic branch, and stratification with respect to these clusters should ensure that each branch is well-represented in both the training and testing data. The cluster assignments of mass spectra thus serve as a proxy for the unknown phylogenetic information. Our hypothesis that clustering the samples into subgroups could reflect true phylogenetic information is motivated by the presence of known phylogenetic lineages within a studied microbial species. For *E. coli* between six and fourteen phylogenetic subgroups are established, depending on the level of granularity [Abram et al., 2019, Clermont et al., 2012]. In *K. pneumoniae*, several subspecies have been characterized, such as *K. pneumoniae (sensu stricto), K. variicola, K. africana*, and other species [Lam et al., 2018a,b, Wick et al., 2018, Wyres et al., 2016]. In *S. aureus*, the presence of sublineages [Bowers et al., 2018] have been described, as well as different taxonomical classes, such as *S. aureus (sensu stricto), S. schweitzeri*, and *S. argenteus* [Tong et al., 2015].

Our contributions. We hypothesise that extending the common approach of stratification by prediction class with stratified cluster labels in train and test will lead to a better and more stable predictive performance. We propose to choose the optimal cluster parameters through clustering validity indices, which are purposefully not using any resistance label information. In this paper, we demonstrate a simple approach to obtain such a stratified assignment using hierarchical clustering on the mass spectra. Our approach is easy to implement, and we demonstrate that it has the ability to improve performance in seven out of nine scenarios for both classification models. Specifically, we observe increases in average precision for ciprofloxacin resistance prediction from 42.3 to 47.1 in *E. coli* and from 44.6 to 50.8 for amoxicillin-clavulanic acid resistance prediction in *S. aureus*.

MALDI-TOF MS. MALDI-TOF MS is an analytical technique, in which a laser decomposes and ionises biomolecular samples into charged molecules, and then determines the mass-to-charge ratio of the ions [Hillenkamp et al., 1991, Proteomics, 2020]. The output is a MALDI-TOF mass spectrum depicting the measured particle intensity against its mass-to-charge ratio. To reduce noise and accentuate peaks, one MALDI-TOF measurement consists of repeated measurements, which are merged into a single output spectrum. Raw spectra suffer from a varying baseline, caused by the matrix solution, and slightly varying peak positions, thus considerably increasing the noise level. To employ such spectra in practice, one hence applies a collection of pre-processing steps, which can be performed using either commercial [Bruker Daltonics, 2018] or open-source software [Gibb and Strimmer, 2012].

## 2 Related Work

MALDI-TOF MS based machine learning is an active and rapidly expanding field of research. While MALDI-TOF MS is already the established method to identify the species causing an infection, several research directions aim to exploit the information contained in MALDI-TOF mass spectra for a more fine-grained characterisation of microbes. We briefly discuss several current research topics.

The most established application for machine learning on MALDI-TOF mass spectra is the identification of sub-species of a probe [Chung et al., 2019, Sonthayanon et al., 2019]. Some phylogenetic lineages within a microbial species are known to cause severe infections, necessitating rapid, high-throughput identification methods. Therefore, quick, reliable and low-cost typing methods of sub-species are essential for an effective infectious disease control. Applying suitable data analysis tools and machine learning to MALDI-TOF mass spectra can provide such a method and is generally of lower cost than current identification methods such as multi-locus sequence typing (MLST) [Chung et al., 2019, Sonthayanon et al., 2019]. Previous work employing MALDI-TOF MS based machine learning for sub-species discrimination includes the typing of *Mycoplasma pneumoniae* [Xiao et al., 2014], discrimination of environmental and contagious *Streptococcus uberis* [Esener et al., 2018], and strain typing *Staphylococcus haemolyticus* [Chung et al., 2019]. Additionally, MALDI-TOF MS has been shown useful for a less expensive, time-consuming and labour-intensive identification of clonal complexes, such as methicillin-resistant *Staphylococcus aureus* (MRSA), vancomycin-intermediately resistant *Staphylococcus aureus* (VISA), and heterogeneous VISA (hVISA) [Camoez et al., 2016, Zhang et al., 2015].

More recently, MALDI-TOF MS based machine learning has been applied for the prediction of antimicrobial resistance [Weis et al., 2020b]. The current standard of culture-based antimicrobial susceptibility testing in clinical routines relies on phenotypic assay approaches, which can take up to four days from sample collection during which broad-spectrum antibiotics are administered [Weis et al., 2022]. Such less specific treatments can cause more serious adverse effects and may ultimately lead to further antibiotic resistances. MALDI-TOF MS based antimicrobial resistance prediction promises to reduce the time required to determine suitable treatment by 48 h to 72 h and minimise the use of broad-spectrum antibiotics [Weis et al., 2022]. Antimicrobial resistance prediction based on MALDI-TOF mass spectra has been proven possible in several scenarios, including carbapenem resistance in *Klebsiella pneumoniae* [Huang et al., 2020], intermediate resistance to vancomycin in *Staphylococcus aureus* [Wang et al., 2018], and carbapenem resistance in *Bacteroides fragilis* [Ho et al., 2017]. This line of research is actively expanding, with large clinical datasets providing information on a variety of species and antimicrobial resistance profiles made publicly available [Weis et al., 2021].

## 3 Materials and Methods

The bacterial species we selected to focus on for this paper are *Staphylococcus aureus* (*S. aureus*), *Escherichia coli* (*E. coli*) and *Klebsiella pneumoniae* (*K. pneumoniae*); these three species were found to be the leading pathogens for deaths associated with antimicrobial resistance [Murray et al., 2022]. Subsequently, we describe all details pertaining to the dataset, including its provenance, its peak calling, and how we use it in a classification scenario.

The data used in this project is a subset of the *DRIAMS* database [Weis et al., 2021]. *DRIAMS* is a large-scale, publicly-available, high quality collection of bacterial and fungal MALDI-TOF mass spectra derived from routinely-acquired clinical isolates [Weis et al., 2022]. To ensure reasonable training times and to allow for comparability with Weis et al. [2020a], we focused on data from the *DRIAMS-A* site collected in 2017 and 2018. Peak-calling was performed with the MALDIquant package [Gibb and Strimmer, 2012] as follows: (i) peak detection using the Median Absolute Deviation (MAD) method with a half-window size of 20 and a signal-to-noise ratio of 2, (ii) defining peaks which, with a tolerance of 400 ppm, appear in at least 90% percent of all spectra as *reference peaks*, and (iii) warping spectra and peaks along the *m*/*z* axis according to linear warping functions determined using the reference peaks and a tolerance of 200 ppm. Please refer to the original dataset for more information on data pre-processing. The resulting dataset is described in Table 1 and includes an average of 218 peaks per MALDI-TOF mass spectrum.

**Table 1:** Summary statistics of the DRIAMS-A [Weis et al., 2021] dataset, considering data collected in 2017 and 2018

| species | antibiotic | scenario | # samples | % prevalence |
| --- | --- | --- | --- | --- |
| *E. coli* | amoxicillin / clavulanic acid | E-AMOXCLAV | 3039 | 20.7 |
|  | ceftriaxone | E-CEF | 3093 | 11.1 |
|  | ciprofloxacin | E-CIPRO | 3081 | 22.8 |
| *K. pneumoniae* | ceftriaxone | K-CEF | 1885 | 6.8 |
|  | ciprofloxacin | K-CIPRO | 1876 | 14.3 |
|  | piperacillin / tazobactam | K-PIPTAZO | 1826 | 9.4 |
| *S. aureus* | amoxicillin / clavulanic acid | S-AMOXCLAV | 2317 | 9.7 |
|  | ciprofloxacin | S-CIPRO | 2408 | 14.9 |
|  | penicillin | S-PEN | 2244 | 70.5 |

Labels and sampling procedure. The antimicrobial resistance labels in Weis et al. [2021] are defined as binary classification problem, with the three susceptibility classes categorised into a positive class, comprising resistant and intermediate samples, and a negative class, comprising all susceptible samples. This label generation process does not always lead to a balanced dataset. Since imbalanced class labels may pose an issue for some machine learning algorithms, we decided to “oversample” the minority class [Lemaître et al., 2017]. The effects of this procedure are most pronounced for the Gaussian Process classifier and less so for logistic regression because this algorithm supports class weights [Weis et al., 2020a].

Sample removal. While the dataset and method closely reflect the setup presented in Weis et al. [2020a], we implemented several changes to impose more quality control steps to the study set-up. First, we removed all *DRIAMS-A* samples collected at the hospital hygiene workstation. At the *DRIAMS-A* collection site, MALDI-TOF mass spectra were collected at different workstations, depending on the type of sample. For example, there are workstations for urine, stool and blood culture samples. One workstation is operated by a hospital’s hygiene department, which is responsible for monitoring of nosocomial infections and the screening for multidrug-resistent pathogens. The growth medium used at this workstation generally contains antibiotics to specifically select for resistant pathogens, which could therefore be reflected in the corresponding MALDI-TOF mass spectrum. To avoid this confounding factor, we chose to remove all samples which were collected at the hospital hygiene workstation. We also improved the stratification process to account for multiple measurements reflecting the same patient case. If multiple measurements originate from the same patient, possibly describing the same infectious strain, it is likely that measurements of these samples show a high degree of similarity. To avoid information leakage between the train and test set, we split such that all samples of the same patient case are either exclusively a part of the train or the test set.

### 3.1 Hierarchical clustering

We use hierarchical clustering to infer a latent phylogenetic structure from MALDI-TOF MS data, which we will subsequently use to improve classification performance. Specifically, we will use the inferred clusters as additional class labels to provide stratification information for splitting the data into a train and test data set, ensuring that the percentage of samples for each joint class–cluster combination are preserved. Stratified train–test splits are necessary to ensure that the testing distribution resembles the training distribution, thus preventing a phenomenon known as “distribution shift”, which typically has adverse effects on generalisation performance. Hierarchical clustering requires either a distance matrix between samples or a feature matrix representation. We opt for the latter and use the data in the same fashion as for the classification, i.e. binned into fixed-size feature vectors. This is a simplification, as it does not account for larger shifts in peak positions, but we find this to be sufficient for our purposes, namely the characterisation of spectra to infer the underlying phylogenetic structure.

#### 3.1.1 Clustering algorithm

We used hierarchical agglomerative clustering [Rokach and Maimon, 2005], which is a bottomup clustering approach: Initially, each sample forms a cluster and the two “closest” clusters— according to a linkage method and a distance measure—are merged into one cluster in each iteration. This process is repeated until only a single cluster remains, containing all samples. Hierarchical clustering is highly flexible: first, a tree summarising the inferred structural relationship between all samples (a *dendrogram*) is constructed, and no *a priori* knowledge about the number of clusters *k* is required; thus, the structure of the dendrogram is not influenced by the choice of *k*, and *k* can be instead selected *a posteriori* using either a threshold on the distances, or it may be inferred from an auxiliary visualisation. A depiction of a dendrogram inferred for the dataset S-AMOXCLAV can be found in Supplementary Figure 1. Moreover, hierarchical clustering is not restricted to specific classes of shapes, as compared to other clustering algorithms like *k*-means.

As the distance between clusters is defined by the selected linkage method and distance metric, both of these components play a crucial role in the clustering process [Nielsen, 2016]. We will subsequently make use of two linkage criteria, *Ward’s linkage* and *average linkage*, always using the Euclidean distance as a ground metric since it is the most common choice for numerical data. We only provide a brief overview here; please refer to the Supplementary Materials for more information on linkage criteria.

Ward’s linkage. Ward’s linkage employs Ward’s minimum variance method [Ward, 1963] and is restricted to the Euclidean distance. Nevertheless, it remains one of the most popular linkage methods.

Average linkage. Average linkage does not only consider particular extremal distances between points, but takes the average distance between all points of two clusters into account. This method is also known as *UPGMA* (unweighted pair group method with arithmetic mean).

#### 3.1.2 Using clusters to improve stratification

Any choice of linkage criterion will lead to a set of clusters containing certain spectra. We hypothesise that clusters are at least implicitly capturing information about phylogeny: if two spectra of the same species are part of the same cluster, we conjecture that they are more related in terms of their phylogeny than they are to any other spectrum from another cluster. Denoting the antimicrobial resistance of a spectrum with *l*_*r*_ and its cluster label with *l*_*c*_, we assign the spectrum a meta-label consisting of (*l*_*r*_, *l*_*c*_) and at the same time keep track of patient case numbers. We consider this meta-label to provide more information about the phylogeny of spectra than the antimicrobial resistance label alone. When splitting a dataset into its train and test parts, we can therefore ensure that the prevalence of each meta-label is equivalent in both parts. Our conjecture is that, assuming the cluster label is implicitly informed by the phylogeny of a spectrum, both parts of the dataset are more similar to each other. This should lead to (i) improved classification performance, as a classifier will encounter similar distributions in the train and test part of the dataset, and (ii) reduced variance between different splits of a dataset, as a classifier will have seen a sufficiently large number of representative samples during training, thus making it possible to generalise better to unseen samples in the test dataset.

Essentially, our method can be seen as providing classifiers with an additional set of inductive biases. Under the assumption that both the train and the test samples arise from the same microbial population, having access to an (estimate of) their phylogenetic information imposes additional structural constraints on downstream models. We find that such constraints improve empirical performance in classification tasks, with the main advantage of our method being the simple integration into any classification workflow. Moreover, our method is easy to implement and, depending on the range of parameters, imposes only minor computational constraints. We merely require the choice of a clustering algorithm (involving a linkage criterion and a distance metric) as well as the choice of the desired number of clusters *k*. The antimicrobial resistance labels are only used for the train–test split; they are *not* required for the clustering step. This makes our method generally applicable; for example, it can be applied to any prediction task and is not restricted to antimicrobial resistance labels.

#### 3.1.3 Clustering output

The cluster algorithm results in a *linkage matrix* that contains information about the distances between individual clusters. Subsequently, we opted for pre-defining the number of clusters *k*, as it allows for an intuitive interpretation to the clustering in terms of number of phylogenetic branches found. Note that this is only required in order to simplify our stratified train–test split; it would also be possible to generate splits for multiple values of *k*, but this goes at the expense of computational performance. We stress that *k* only constitutes an upper bound of the number of clusters; depending on the linkage criterion, it is possible that fewer than *k* cluster can be formed. For instance, in some of our experiments for *E. coli*, we were unable to obtain *k* = 2 clusters when using single linkage. This was caused by three clusters having the same linkage distance, thus resulting directly in a merge into one larger cluster.

#### 3.1.4 Clustering metrics

The main challenge when applying clustering algorithms is to pick a suitable number of clusters *k*. In many applications—including ours—the “correct” label (here, the phylogenetic structure of a sample) is unknown *a priori*. To this end, numerous *clustering validity measures* were developed, which help assess the quality of a clustering in the absence of ground truth labels. We selected clustering validity metrics that were shown to perform well, provided the feature space does not give rise to highly-complex topological structures [Rieck and Leitte, 2016] such as cycles or voids. Notice that the clustering validity metrics that we used for this study are *unsupervised* and do *not* have access to the antimicrobial class labels.

Silhouette score. The Silhouette score takes into account the mean distance between a point and all other points belonging to the same cluster (denoted as *a*), and the mean distance between a point and all points in the nearest cluster (denoted as *b*). The Silhouette coefficient for a single point is then defined as *s* = *b*−*a*/max(*a,b*) and the mean of all local Silhouette coefficients gives rise to the overall score. We have *s* ∈ [−1, 1], with scores close to −1 indicating a bad clustering, and scores close to 1 indicating highly dense and well-separated clusters.

Davies–Bouldin index. The Davies–Bouldin index indicates the average similarity between each cluster and its most similar cluster, where similarity is defined as the ratio of within-cluster distances to between-cluster distances. Let *s*_*i*_ be the cluster diameter, i.e. the average distance between each point of the cluster and its corresponding centroid, and let *d*_*ij*_ be the distance between the centroids of cluster *i* and *j*. The Davies–Bouldin index is then defined as 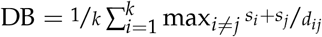, where *k* is the number of clusters. The Davies–Bouldin index is always non-negative, with *lower* values signifying a better clustering and 0 being the best possible score.

### 3.2 Classification

Following Weis et al. [2020a], we used a logistic regression classifier (LR) and a Gaussian Process classifier based on the Peak Information Kernel (GP-PIKE), which was specifically designed to handle sparse MALDI-TOF mass spectra.

Logistic regression. Logistic regression requires inputs of a fixed length. Therefore, the classification pipeline for this model contains a binning step that ensures that all spectra are represented by fixed-length feature vectors. Since information about the number of bins is usually not available, we consider the number of bins to be a hyperparameter of the model that we will tune using a grid of {300, 600, 1800, 3600} bins. Additionally, following Weis et al. [2020a], different choices of regularisation (*L*_1_, *L*_2_, *elastic net*, and no regularisation) as well as the regularisation strength parameter *C* ∈ {10^−4^, 10^−3^, …, 10^3^, 10^4^} are optimised using cross-validation. We recorded the optimal hyperparameters for each experiment and generally observed considerable variations in model selection for different species–antibiotic combinations.

Gaussian Process with Peak Information Kernel. The GP-PIKE classifier is designed to work with sparse MALDI-TOF MS peaks, provided in the form of sets of tuples [Weis et al., 2020a]. PIKE is a kernel inspired by heat diffusion processes, capable of capturing (non-linear) interactions between individual peaks. The kernel does not require binning and can make direct use of the peaks. The integration of PIKE into a Gaussian Process classifier has two primary advantages: first, the kernel hyperparameter can be optimised using maximum likelihood; it is therefore *not* restricted to a predefined parameter grid (model averaging is also simplified, as the smoothing parameters learnt for different runs can be harmonised by calculating their mean). Second, GP-PIKE returns confidence estimates and offers the ability to *reject* the classification of out-of-distribution samples. This capability is crucial in clinical settings, as clinicians need to be able to rely on decisions made by a classifier [Weis et al., 2020a].

Performance evaluation. We used five random seeds to split the data and for each of these splits, we employed a 5-fold cross-validation procedure with stratified folds to optimise hyperparameters based on average precision. Average precision approximates the area under the precision–recall curve (AUPRC) and is a suitable performance metric when working with heavily-imbalanced classes.

## 4 Results

Inferring hierarchical structure. We clustered the datasets for each species five times, using different linkage criteria (*ward, average, weighted, single, complete*). The number of clusters were varied from *k* = 1 (no clustering) to *k* = 20, which we consider to be a biologically plausible range. An illustration of the hierarchical clustering on the S-AMOXCLAV dataset using *Ward’s* linkage criterion is depicted in Supplementary Figure 1. For each clustering, we report two unsupervised clustering validity metrics, the *Silhouette score* and the *Davies–Bouldin index*, to judge how well structures in the data were separated. We once again stress that we do not have ground-truth strain-type labels available, hence our need for unsupervised validity metrics. Figure 1 depicts the behaviour of the cluster validity metrics with respect to varying the number of clusters *k* for *average* linkage. The validity metrics for the remaining linkage methods can be found in Supplementary Figure 3.

**Figure 1:**
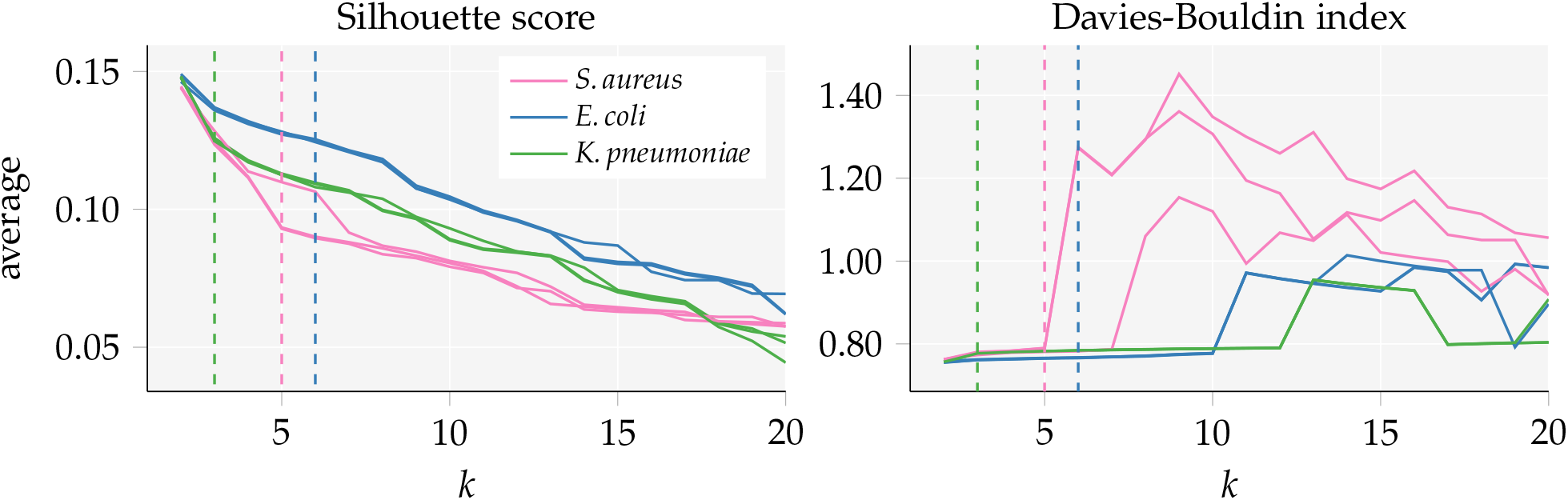
Behaviour of the Silhouette coefficients and Davies-Bouldin index using the average linkage criterion with varying number of clusters (k ≥ 2 because all scores require at least two clusters). A Silhouette score close to 1 indicates well separated clusters and values less then 0 indicate that a number of samples were assigned to the wrong cluster; a low Davies-Bouldin index indicates good separation, the clusters are far apart and show little dispersion. Three curves are depicted per species (one curve per species-antibiotic dataset, each with different samples). The chosen number of clusters k^∗^ is indicated by a dashed line for each species.

We report three curves per species, as each species dataset contains slightly different samples depending on which antibiotic is considered; please refer to Table 1 for more details. In general, we observe that the best values (in terms of the clustering validity index) are obtained for the smallest number of clusters, i.e. *k* = 2.

Choosing the number of clusters. We have to select a *single* clustering in order to obtain a set of labels that we can incorporate into a stratified train–test split. This requires determining an optimal value for the parameter *k*, which we will refer to as *k*^∗^. This choice should be data-driven, without taking any information about antimicrobial resistance or predictive performance into account. We therefore base this choice solely on clustering validity indices depicted in Figure 3, obtaining a fixed *k*^∗^ for each species, which we will subsequently use for the stratified train–test split. Specifically, we analysed local maxima in the *Silhouette score* and local minima in the *Davies-Bouldin index*, picking values for *k*^∗^ in their immediate vicinity. Additionally, we considered values for *k*^∗^ for which the curves exhibit an elbow or around which we observed a rapid change in the clustering validity index. We thus derived the final values as *k*^∗^ = 6 for *E. coli, k*^∗^ = 3 for *K. pneumoniae* and *k*^∗^ = 5 for *S. aureus*. Subsequently, we will use the *ward* and *average* linkage criteria, as they are the most common choice for data analysis tasks.

For *E. coli* we see a drop in the silhouette coefficient value for *k* > 6, while the Davies–Bouldin index in many cases seems to dip for *k* = 6 (recall that this index should be minimised, whereas the silhouette score should be maximised). For *K. pneumoniae* we observe such a drop in the silhouette score at *k* = 3 and a steep increase in the Davies–Bouldin index for *k* > 3 (see *ward* and *complete* linkage in supplementary material). A similar increase in the Davies–Bouldin index can be observed for *k* > 5 for *S. aureus*.

Resistance prediction. Using the optimal choice of the number of clusters *k*^∗^, we use the resulting clusters to stratify the train–test splits, serving as additional class labels, next to the labels that we defined based on the antimicrobial resistances profiles and taking into account that samples coming from the same patient should all be either in the training or testing set. We then incorporate these train–test splits into two antimicrobial resistance classification scenarios; one using a logistic regression classifier and one using GP-PIKE, the Gaussian process (GP) in combination with the Peak Information Kernel (PIKE), specifically designed for MALDI-TOF mass spectra [Weis et al., 2020a]. For each of the experiments, we report the mean and standard deviation of the average precision score over the random seeds. Due to computational constraints, it was only possible to evaluate the GP-PIKE model in four of the five seeds in some experiments, and in a handful of cases, the results were calculated over three seeds.

The prediction results for both classifiers with and without using the hierarchical stratification are depicted in Table 2. We focus on two linkage criteria, namely *ward* and *average*. This was motivated by the fact that *ward* remains a popular choice, at least when the Euclidean distance is selected as a ground metric. In addition, we selected the *average* linkage criterion as a trade-off between the *single* linkage criterion (with known issues such as chaining) and the *complete* linkage criterion (which is computationally infeasible for large-scale datasets).

**Table 2:** Improved performance of train-test splits including clustering information compared to a standard random train-test split. Results of three methods given by mean AUPRC ± standard deviation on all random splits. We have k^∗^ = 6 for E. coli, k^∗^ = 3 for K. pneumoniae and k^∗^ = 5 for S. aureus.

| scenario | LR<br>$k = 1$ | LR<br>$k^*$ (ours)<br>ward | LR<br>$k^*$ (ours)<br>average | GP-PIKE<br>$k = 1$ | GP-PIKE<br>$k^*$ (ours)<br>ward | GP-PIKE<br>$k^*$ (ours)<br>average |
| --- | --- | --- | --- | --- | --- | --- |
| E-AMOXCLAV | 24.5±1.4 | 25.7±3.7 | **25.8±2.4** | 24.9±2.5 | **27.7±2.8** | 26.0±2.7 |
| E-CEF | 19.4±5.4 | **22.9±7.0** | 21.1±4.2 | **20.9±6.4** | 20.6±6.0 | 19.9±7.5 |
| E-CIPRO | 38.8±4.7 | 39.9±3.5 | **40.5±3.7** | 42.3±6.9 | 44.3±4.6 | **47.1±3.8** |
| K-CEF | 16.5±2.3 | 16.2±9.4 | **17.6±4.7** | **28.8±12.2** | 20.1±15.3 | 27.4±14.9 |
| K-CIPRO | 26.5±3.7 | 26.4±6.6 | **27.9±4.7** | 29.9±5.4 | 29.8.2±8.4 | **31.1±6.6** |
| K-PIPTAZO | 14.1±2.8 | 14.0±2.7 | **15.6±5.0** | 16.5±4.3 | **18.9±7.2** | 17.6±5.4 |
| S-AMOXCLAV | **28.4±4.4** | **28.4±2.2** | 27.2±4.8 | 44.6±3.1 | **50.8±5.8** | 47.5±6.1 |
| S-CIPRO | **30.7±5.3** | 26.5±10.3 | 30.3±4.4 | 35.4±6.4 | 32.0±10.2 | **40.6±5.0** |
| S-PEN | 75.4±1.8 | 76.2±4.2 | **77.2±3.4** | 79.9±1.7 | 80.7±3.0 | **81.0±2.4** |

For both classification models, seven out of the nine prediction tasks exhibit performance improvements using our novel proposed hierarchical stratification. In logistic regression, improvements are highest for ceftriaxone resistance prediction in *E. coli*, with an average precision increase from 38.8 to 40.5. GP-PIKE started at a higher baseline performance, with increases in predictive performance from 44.6 to 50.8 for amoxicillin-clavulanic acid resistance prediction in *S. aureus*.

For the sake of understanding the relationship between *k*, clustering validity values, and predictive performance, we depict the predictive performance for all *k* ∈ {1, 2, …, 20} in the supplementary material; see Supplementary Figure 4 for logistic regression and Supplementary Figure 5 for GP-PIKE. We emphasise the need to fix *k prior* to considering the average precision results, as was done for the choice of *k*^∗^ in this publication, in order to not let information leakage influence the choice of the clustering parameters. We observe a high sensitivity to the stratification for predicting antimicrobial resistance to all three antibiotics for *K. pneumoniae*, and also for amoxicillin-clavulanic acid resistance in *S. aureus*. Our results therefore do not support the hypothesis that hierarchical stratification leads to a lower standard deviation. We observe, however, that hierarchical stratification can improve predictive performance. It becomes evident that the choice of *k strongly* influences whether the stratification will increase the predictive performance. In many scenarios, a poor choice of *k* led to a decreased performance, as compared to the baseline of no clustering, *k* = 1.

## 5 Discussion

We presented a novel stratification for MALDI-TOF MS based phenotype classification tasks, based on hierarchical structures inferred from the MALDI-TOF mass spectra. Our novel proposed stratification was driven by the hypothesis that clusters would be able to implicitly capture phylogenetic structure. This required the use of clustering validity scores in order to choose the number of clusters *k*. Our experiments indicated that the clustering validity scores *Silhouette score* and *Davies-Bouldin index* did not give a clear picture as to which value of *k* would be an optimal choice to obtain well-separated clusters. We nevertheless demonstrated the general utility of learning such stratifications to improve predictive performance in the context of antimicrobial resistance prediction.

We observed that the stratification can influence predictive performance quite drastically (despite the fact that for every choice of *k*, the same number of samples are being used for training, albeit with a different arrangement in the train and test part, respectively). Good performance— and improved performance compared to considering no hierarchical stratification—likely stems from the fact that both train and test dataset follow the “true” structure of the data closely. Therefore, each split (induced by each seed) is capable of training on a dataset that closely follows the true distribution.

Despite the observed performance improvements, we also found hierarchical stratification to be incapable of reducing the standard deviation in the prediction results. This could be due to the hierarchical clustering not capturing the structure that caused this high standard deviation. It could also be caused by stratifications that do not account for the complexity of classifying certain parts of the data, leading to an under- or overestimation of predictive performance. An alternative hypothesis would be that the high variance in predictive performance stems from the small samples sizes, as the species with the smallest sample size, *K. pneumoniae*, exhibited a larger standard deviation than the other two species. Overall, we consider these results to be encouraging; they highlight the potential of including phylogenetic information, but also indicate that this endeavour is fraught with difficulties that need to be overcome in order to yield a stable method, namely (i) no ground truth to validate the clustering, and (ii) obtaining a good choice for *k*^∗^, the optimal number of clusters, that is beneficial for *all* prediction tasks.

Our proposed stratification method is but a first step in this direction—while it is conceptually simple and easy to implement, increasing the stability of the results will require additional research. We thus foresee multiple future strands of research. First, collecting a dataset that includes strain-type information on the bacterial species would allow to quickly validate if the clustering is in fact capturing the phylogenetic relatedness of samples or not. Detailed information on additional latent information, such as the growth medium used for culturing, reflected in the MALDI-TOF mass spectra might provide further insights into confounding factors influencing the clustering results. Second, we conjecture that the results are highly influenced by the choice of metric for the clustering. We restricted our experiments to the Euclidean distance for conceptual simplicity, but the development of domain-specific metrics bears the promise of being able to capture multi-scale differences between spectra while being impervious to noise. We are particularly interested in methods based on *optimal transport* [Villani, 2009], which have shown promising performance recently in classification tasks. Finally, it would also be interesting to develop a fully-automated way of choosing k^∗^ in practice; this choice could potentially be driven by more advanced clustering validity indices.

## Author Contributions

C.W., B.R. designed machine learning experiments; C.W., B.R., S.B. implemented all experiments of the machine learning analysis; A.C., A.E. organised collection and extracted clinical data; A.C, C.W. implemented pre-processing of the dataset; C.W., B.R., S.B. wrote the manuscript with the assistance and feedback of all co-authors; K.B. supervised the study.

## Supplementary Material

### Linkage criteria

Weighted linkage. The weighted linkage criterion considers the weighted mean of average distances between cluster members. Specifically, the distance between cluster *u*, which was formed by merging clusters *s* and *t*, and any other cluster *v* is defined as

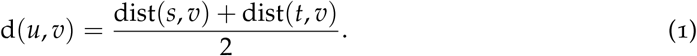

The initial distances between singleton clusters are given by the selected distance metric, i.e. for singleton clusters *a* and *b*, we have d(*a, b*) = dist(*a, b*). This process is computationally simpler than average linkage but because of the weighted approach, distances do not contribute equally during calculation. The weighted linkage criterion is also known as *WPGMA* (weighted pair group method with arithmetic mean).

Single linkage. The single linkage criterion is arguably the most efficient implementation for agglomerative hierarchical clustering. It defines distances between clusters *u* and *v* as

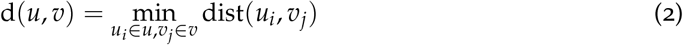

for all members *u*_*i*_, *v*_*j*_ in cluster *u* and *v*, respectively. As single linkage method only considers the *minimum* distance, they suffer from a phenomenon known as *chaining*, whereby cluster shapes tend to form “chains” in the feature space, leading to heavily-imbalanced cluster sizes. It its therefore possible for two clusters to be considered very close even though the majority of their points are very distant, only because of a small subset of outliers that are close to each other but assigned to different clusters.

Complete linkage. The aforementioned disadvantage of single linkage can be avoided by the *complete* linkage method. It examines the *maximum* distance between points of two clusters, i.e.

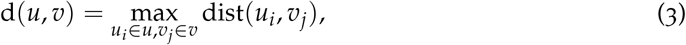

where *u*_*i*_, *v*_*j*_ refer to points in cluster *u* and cluster *v*, respectively. Similar to single linkage, this criterion is *also* sensitive to outliers.

### Inferring hierarchical structure

**Supplementary Fig. 1:**
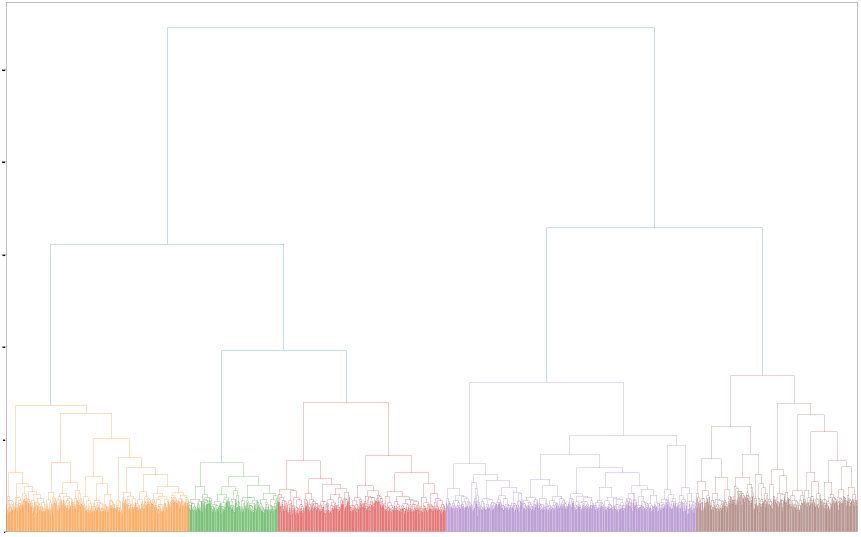
Dendrogram of the complete hierarchical clustering tree of all data points in S. aureus and penicillin using Ward’s linkage. Each tip at the lower end of the tree corresponds to one sample in the dataset. The clusters derived from the chosen k^∗^ = 5 are indicated by colors. Note that the clustering is independent from the number of clusters; the cluster labels are only derived after the dendrogram is defined through a cut-off at the desired “height” of the tree.

**Supplementary Fig. 2:**
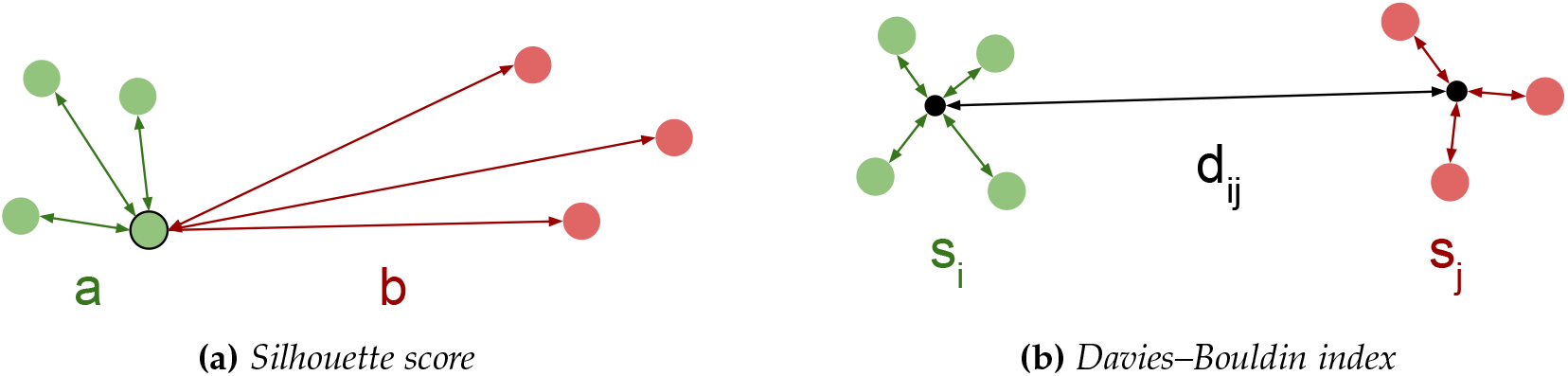
A depiction of the clustering validity scores’ functionality: (2a) The average distance of one point to all points belonging to the same cluster is denoted as a, while the average distance to the nearest foreign cluster is denoted as b. (2b) The average distance of each point in a cluster to the cluster’s centroid point are denoted as s_i_ and s_j_ for clusters i and j respectively. These average within-cluster distances are compared with the distance between cluster centroids d_ij_.

Supplementary Figure 3 depicts the behaviour of the cluster validity metrics with respect to varying the number of clusters *k* We report three curves per species, as each species dataset contains slightly different samples depending on which antibiotic is considered; please refer to Table 1 in the main document for more details. In general, we observe that the best values (in terms of the clustering validity index) are obtained for the smallest number of clusters, i.e. *k* = 2. We moreover observe that different linkage criteria give rise to markedly different behaviours in the clustering validity scores. For example, the *Silhouette score* values for *single* and *average* linkage are monotonically decreasing, but appear to reach a plateau for *k* > 10 clusters using the *ward* linkage criterion. We also observe that some linkage criteria are highly sensitive to the small differences in the dataset composition for different antibiotics. This effect is most pronounced in the *Davies–Bouldin index* when applying *average* linkage, or in both scores for the *weighted* linkage.

**Supplementary Fig. 3:**
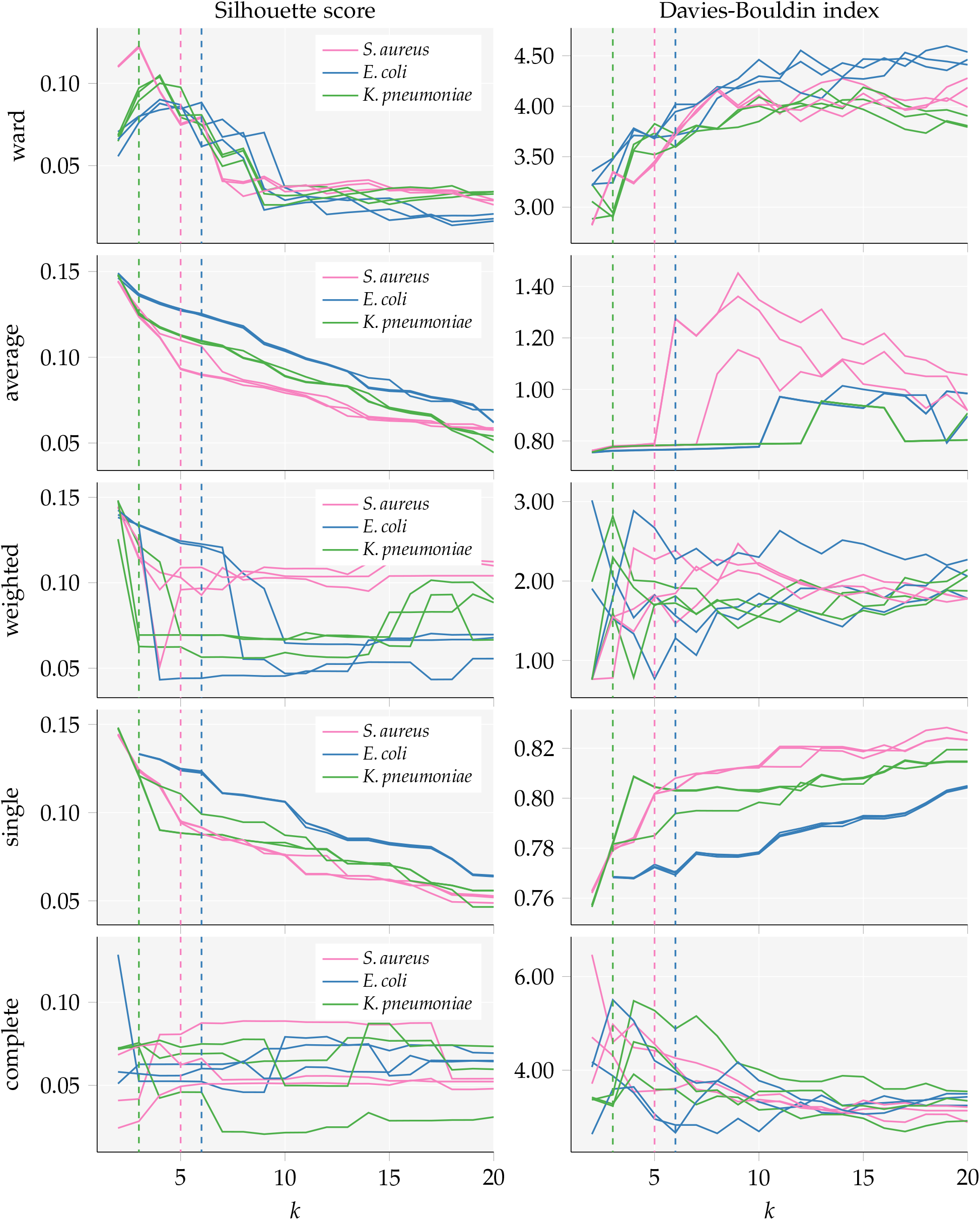
Behaviour of the Silhouette coefficients and Davies-Bouldin index with varying number of clusters. Five different linkage criteria were considered for clustering. Please note that a minimum number of two clusters is needed to compute these clustering scores, and therefore no scores can be reported for k = 1. Please also note the missing validity scores for E. coli using single linkage for k = 2. As described, it is in some cases impossible to exactly form the desired number of clusters. A Silhouette score close to 1 indicates well separated clusters and values less then 0 indicate that a number of samples were assigned to the wrong cluster; a low Davies-Bouldin index indicates good separation, the clusters are far apart and show little dispersion. Three curves are depicted per species, one curve per species-antibiotic dataset, as each dataset contains different samples. The chosen number of cluster k^∗^ is indicated by a dashed line for each species.

**Supplementary Fig. 4:**
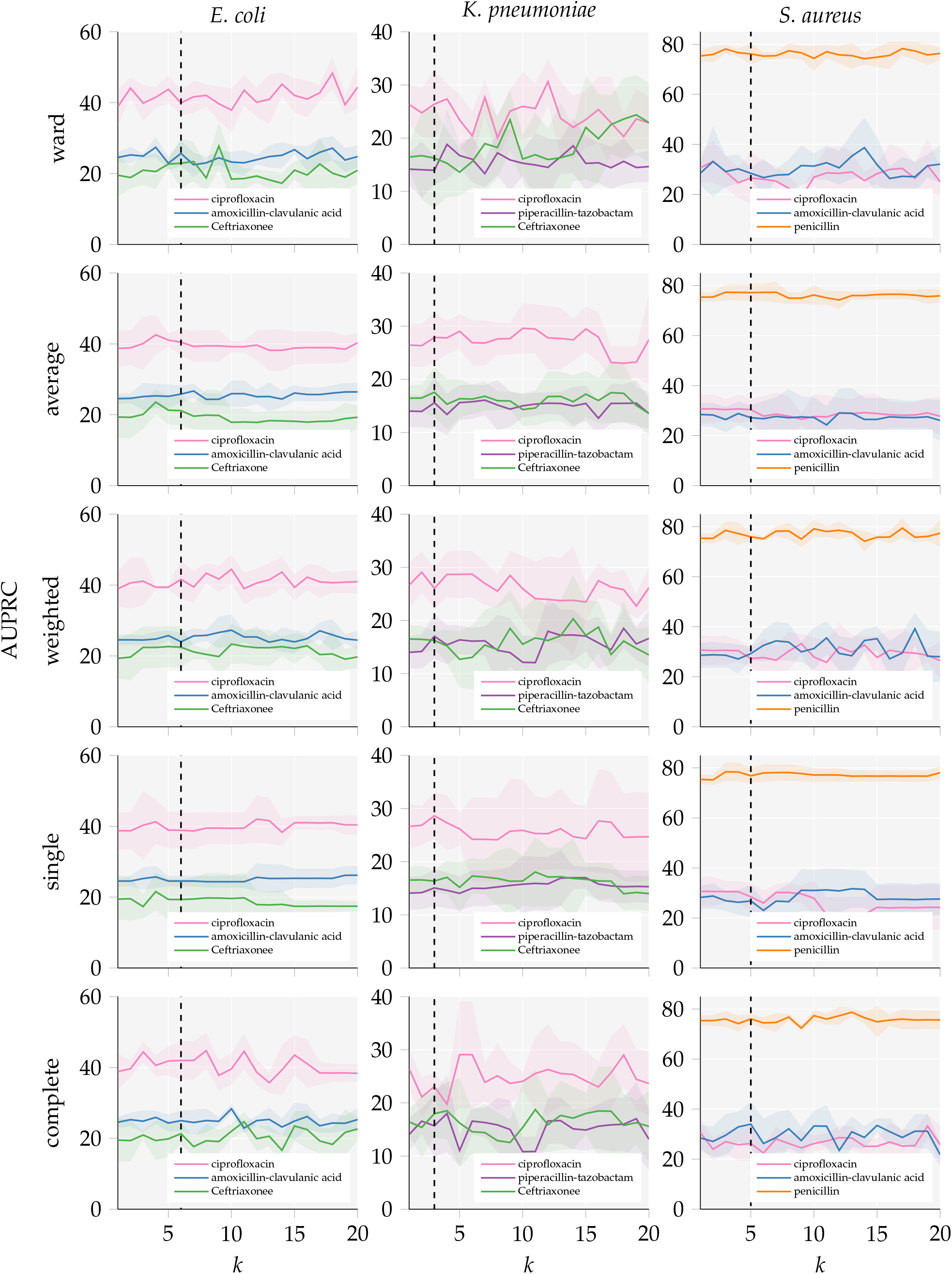
The curves depict the logistic regression prediction performance of all nine antimicrobial resistance scenarios, illustrating the influence of different linkage criteria and the number of clusters k used for hierarchical clustering. All results are given by mean average precision (AUPRC) ± standard deviation on the test fold for all computed random splits. A vertical dashed line indicates k^∗^ chosen for each species.

**Supplementary Fig. 5:**
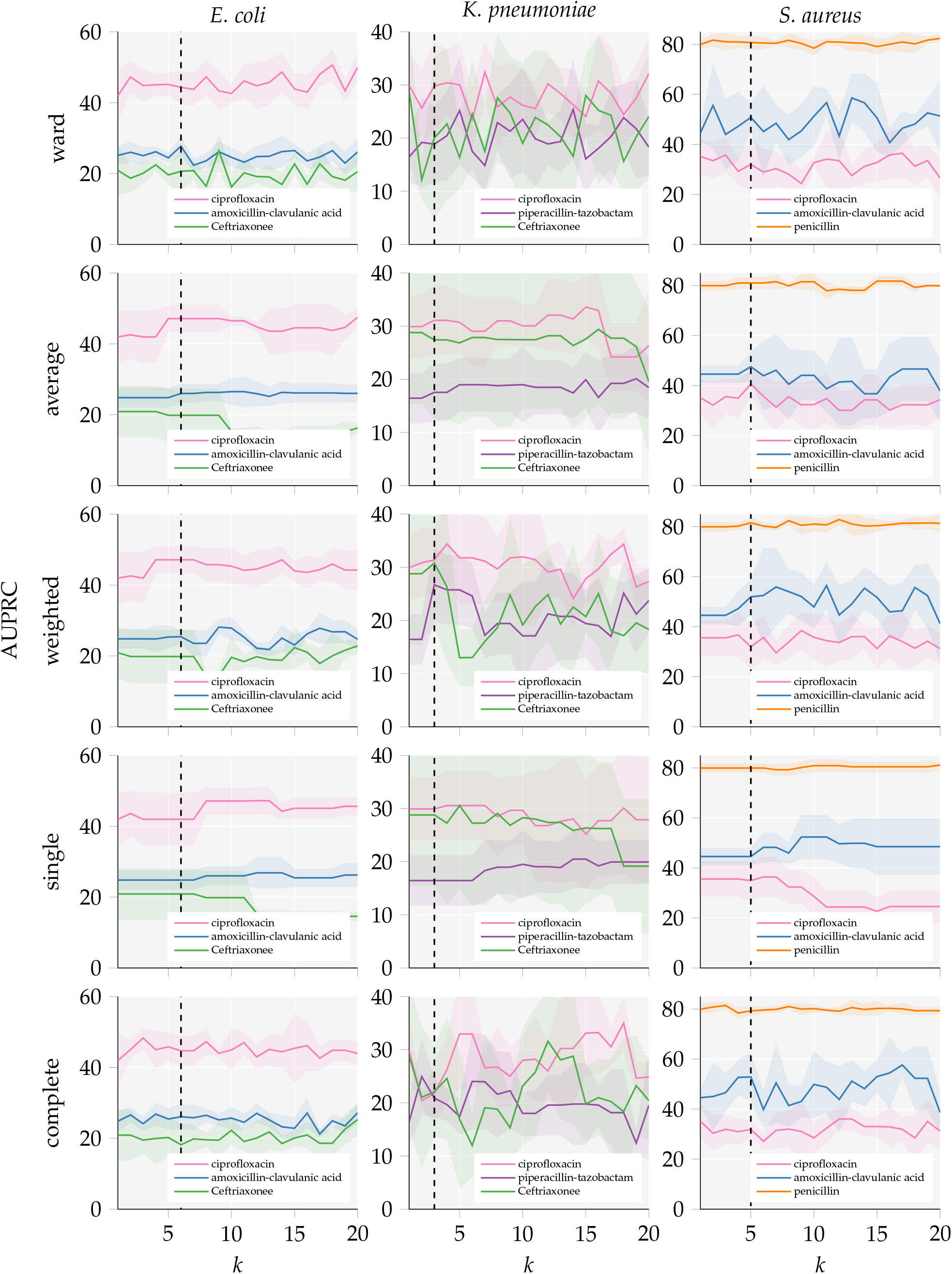
The curves depict the GP-PIKE prediction performance of all nine antimicrobial resistance scenarios, illustrating the influence of different linkage criteria and the number of clusters k used for hierarchical clustering. All results are given by mean average precision (AUPRC) ± standard deviation on the test fold for all computed random splits. A vertical dashed line indicates k^∗^ chosen for each species.

